# The regulation of circadian rhythm by insulin signaling in *Drosophila*

**DOI:** 10.1101/2022.04.25.489482

**Authors:** Sho T Yamaguchi, Jun Tomita, Kazuhiko Kume

## Abstract

Circadian rhythm is well conserved across species and relates to numerous biological functions. Circadian misalignment impairs metabolic function. Insulin signaling is a key modulator of metabolism in the fruit fly as well as mammals and its defects cause metabolic disease. Daily diet timing affects both circadian rhythmicities of behavior and metabolism. However, the relationship between circadian clock and insulin signaling is still elusive. Here, we report that insulin signaling regulates circadian rhythm in *Drosophila melanogaster*. We found the insulin receptor substrate mutant, *chico*^*1*^, showed a shorter free-running circadian period. The knockdown of *insulin receptor* (*InR*), or another signaling molecule downstream of *InR, dp110*, or the expression of a dominant-negative form of InR resulted in the shortening of the circadian period and diminished its amplitude. The impairment of insulin signaling both in all neurons and restricted circadian clock neurons altered circadian period length, indicating the insulin signaling plays a role in the regulation of circadian rhythm in clock cells. Among 3 insulin-like ligands expressed in the brain, *dilp5* showed the largest effect on circadian phenotype when deleted. These results suggested that insulin signaling contributes to the robustness of the circadian oscillation and coordinates metabolism and behavior.

**Highlights:**

- Insulin receptor substrate mutant, *chico*^*1*^, displayed circadian rhythm phenotype.
- Pan-neuronal inhibition of insulin receptor signaling shortened circadian cycle.
- Inhibition of insulin signaling only in clock neurons altered circadian cycle.
- Dilp5 is a major insulin receptor ligand for circadian effects.

## Introduction

Intrinsic circadian rhythms exist in many species and are involved in a variety of physiological phenomena (Dunlap, 1999; Green et al., 2008; Roenneberg and Merrow, 2005). In humans, circadian misalignment has been shown to cause changes in gene expression as well as affect metabolic functions such as plasma insulin and glucose (Archer et al., 2014; Scheer et al., 2009). It has been suggested that such changes due to disruption of circadian rhythm increase the risk of obesity and diabetes (Depner et al., 2014; Noh, 2020).

The fruit fly, *Drosophila melanogaster*, has been used as a genetic model organism to reveal genes regulating circadian rhythm and sleep (Bedont et al., 2021; Dubowy and Sehgal, 2017; Toda et al., 2019; Tomita et al., 2017). We pioneered its sleep regulation (Kume et al., 2005; Tomita et al., 2015, 2011; Ueno et al., 2012; Ueno and Kume, 2014). To date, we examined the relationship between sleep and metabolism or feeding behavior and found that c-Jun N terminal kinase, Sik3, and nutrition or taste of the food function to regulate sleep (Funato et al., 2016; Hasegawa et al., 2017; Takahama et al., 2012; Yamazaki et al., 2012).

Metabolism and feeding behavior tightly relate to not only sleep but also circadian rhythm. For example, null mutation of clock genes disrupts glucose homeostasis and lipid metabolism as in mammals (Rudic et al., 2004; Seay and Thummel, 2011; Turek et al., 2005). Furthermore, feeding with a high-fat diet or high sucrose diet affected clock gene expression or daily activity profile (Lee et al., 2021; Nayak and Mishra, 2021). However, the molecular mechanisms of how metabolism affects circadian rhythm are still elusive.

Insulin signaling, a critical regulator of metabolism in mammals, is well conserved in flies (Nässel et al., 2015; Nässel and Broeck, 2016). Flies have eight mammalian insulin homologs, *drosophila insulin*-*like peptides* (*dilps*), one *insulin receptor* (*InR*), and one mammalian insulin receptor substrate homolog, *chico*. These molecules have been shown to regulate not only metabolism (Rajan and Perrimon, 2012; Ugrankar et al., 2015) and development (Böhni et al., 1999; Brogiolo et al., 2001; Fernandez et al., 1995; Sano et al., 2015) but also behaviors, such as memory, feeding behavior, courtship, and sleep (Barber et al., 2021; Metaxakis et al., 2014; Naganos et al., 2016; Tanabe et al., 2017; Watanabe and Sakai, 2016). In addition to these reports, previous research has shown that *Akt* and *target of rapamycin* (*TOR*), downstream components of insulin signaling, affect circadian rhythm (Zheng and Sehgal, 2010). Although the result suggested that insulin signaling regulates circadian rhythm, it is not clear whether *InR* and *chico* in the central nervous system regulate circadian rhythm. In this study, we aimed to clarify the regulation of circadian rhythm by neuronal insulin signaling.

## Material & Methods

### Fly strains and culture conditions

Flies (*Drosophila melanogaster*) were reared with conventional food, which contained yeast, glucose, cornmeal, wheat germ, and agar, at 24.5°C under a 12-h:12-h light: dark (LD) cycle as described before (Kume et al., 2005). The following stocks were obtained from the Bloomington Stock Center, Indiana University: *elav*-Gal4 (Stock number: 458), *tim*-Gal4 (7126), UAS-*InR* DN (8252), *chico*^*1*^ (10738), *dilp2* KO (30881), *dilp3* KO (30882), *dilp5* KO (30884), *dilp2, 3*, and *5* KO (30889), *y*^*1*^ *v*^*1*^; *P{CaryP}attP2* (36303), *y*^*1*^ *v*^*1*^; *P{CaryP}attP40* (36304), UAS-*chico* RNAi (36665), UAS-*InR* RNAi (51518), and UAS-*dp110* RNAi (61182). *DvPdf*-Gal4 (Bahn et al., 2009) and *Pdf*-Gal4 was provided by Dr. Taishi Yoshii. *Pdf*-GS was gifted from Dr. Takaomi Sakai. UAS-*dilp5* RNAi (VDRC 49520), UAS-*Dicer-2* (60008), and *w*^*1118*^ (60000) which is the genetic background of the UAS-*dilp5* RNAi line were purchased from Vienna Drosophila RNAi Center (VDRC) (Dietzl et al., 2007). *elav*-Gal4, *tim*-Gal4, *dilp2* KO, *dilp3* KO, *dilp5* KO, and UAS-*InR* DN were backcrossed to the control strain (*w*^*1118*^) at least five times.

### Circadian behavioral analysis

The two-to five-days-old male flies were loaded into a glass tube (length, 65 mm; inside diameter, 3 mm) with sucrose food (1% agar supplemented with 5% sucrose) at one end and were entrained for at least 3 days under LD cycle conditions before being transferred to constant dark (DD) conditions. In the experiment with *Pdf*-GS, the two-to five-days-old male flies were loaded into a glass tube with sucrose food containing vehicle (EtOH, - RU486) or 100 µM RU486 (+ RU486) at one end and transferred to DD conditions. Data were collected in 1 min bin for 10 days under DD conditions using the *Drosophila* Activity Monitor system (Trikinetics). The period length and amplitude of circadian rhythm were calculated using locomotor activity data for 9 or 10 days by Fourier transform (FT) with a custom-made R code. Flies with the maximal value of the power spectrum calculated by the FT less than 0.04 were considered arrhythmic and were removed from the calculation of the mean of circadian period length. The data from the experiments of the *InR* DN expression with *elav*-Gal4, *tim*-Gal4, and *DvPdf*-Gal4, or the progenies from crosses between *dilps* KO and TKO were also used in the published paper (Yamaguchi et al., 2021). However, the analysis methods are completely different between this study and the published paper.

### mRNA analysis with quantitative RT-PCR

mRNA levels were examined by quantitative RT-PCR. The 2-to 5-days-old male flies were collected at zeitgeber time 6. Fifteen to thirty heads were divided into three and used for total RNA extraction with RNAiso Plus (TAKARA) according to the manufacturer’s instruction. cDNA was synthesized from the total RNA using ReverTra Ace qPCR Master Mix with gDNA Remover (Toyobo), then used for quantitative RT-PCR using THUNDERBIRD SYBR qPCR Master Mix (Toyobo). *GAPDH2* expression levels were used as an internal control. The primers were 5’-AGGAAGGAAAAGCGGAAAAG-3’ and 5’-GGGGAGGATAAACGAGGTGT-3’ for *GAPDH2*, 5’-GCTTCGCTGCATTTCAACTT-3’ and 5’-AGTGTCCCAGGCGCTTATTA-3’ for *chico*, 5’- GGACAAGGAGCGAATCAAAC-3’ and 5’-TGCAGTTCTCCAGCTGATTG-3’ for *InR*, 5’- CCATCCGGTGCTTACACTTT-3’ and 5’-AGATCGCTTTCCAGGTACGA-3’ for *dp110*, 5’- CCAGGAAGGTTGTCATCTCG-3’ and 5’-GAAAGCCCGCTCGTAGATAAG-3’ for *4E-BP*, 5’- TGAGTATGGTGTGCGAGGAG-3’ and 5’-ACAAACTGCAGGGGATTGAG-3’ for *dilp2*, 5’- AGAGAACTTTGGACCCCGTGAA-3’ and 5’-TGAACCGAACTATCACTCAACAGTCT-3’ for *dilp3*, 5’- GAGGCACCTTGGGCCTATTC-3’ and 5’-CATGTGGTGAGATTCGGAGCTA-3’ for *dilp5*. The primer sequences for *dilp3* and *dilp5* are referred from a previous report (Broughton et al., 2005). Each mRNA expression level was normalized to GAPDH2, then normalized to the average of three independent control samples.

### Statistical analysis

Data were analyzed as described in the figure legend using Microsoft Excel and the freely available statistical software package R 3.5.3 (https://www.r-project.org/).

## Results

### *chico*^*1*^ mutation caused a shorter circadian period

To investigate the relationship between insulin signaling and circadian rhythm, we used the genetic mutant related to insulin signaling. Because homozygous *InR* null mutant is lethal, we used *chico*^*1*^, *chico* mutant line which has P element insertion at 80 bp downstream of translation start site and this insertion cause complete loss of chico protein to examine the effect of InR signal on circadian rhythm (Böhni et al., 1999; Naganos et al., 2012). To remove the genetic background effect, we outcrossed *chico*^*1*^ for five generations into *w*^*1118*^, but the homozygous outcrossed *chico*^*1*^ mutant was lethal. Therefore, we measured the activity of the heterozygous mutants of outcrossed or original *chico*^*1*^ with *w*^*1118*^ (*chico*^*1*^ / + (BC5), *chico*^*1*^ / + (F1)) in addition to the homozygous mutants of original *chico*^*1*^ to investigate the effect of *chico*^*1*^ mutation. *chico*^*1*^ showed a shorter circadian period length (mean ± SEM; 1395 ± 4.78) than the other groups (Fig. 1A, B), although the results were not directly comparable due to the difference in genetic background. Nonetheless, there was little difference in circadian period length between the *chico*^*1*^ / + (BC5) (1421 ± 4.05) and *chico*^*1*^ / + (F1) (1420 ± 2.01), implying the little genetic background effect on circadian period length. Moreover, *chico*^*1*^ / + (BC5) tended to have a shorter circadian period length than *w*^*1118*^ (+, 1433 ± 4.95). These results suggested that *chico* relates to the regulation of circadian rhythm and its defect leads to a shorter circadian period.

**Figure 1.**
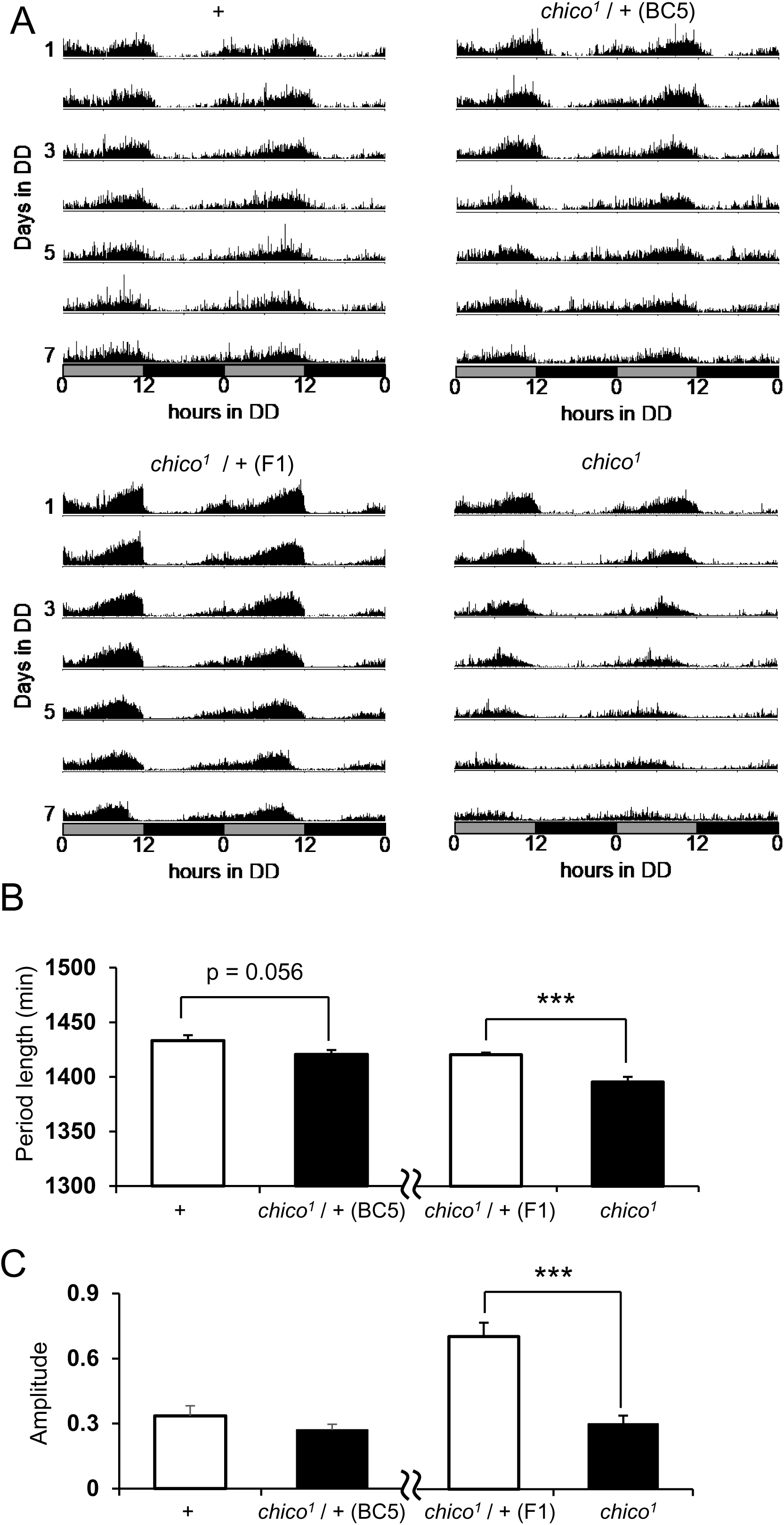
*chico*^*1*^ mutation caused a shorter circadian period length. (A) Average activity from wild type (+, n = 14), backcrossed heterozygous (*chico*^*1*^ / + (BC5), n = 15), original heterozygous (*chico*^*1*^ / + (F1), n = 31), and original homozygous (*chico*^*1*^, n = 30) *chico* mutant flies. Each fly activity was recorded for 10 days in constant dark (DD) conditions. Activity counts over 7 days in DD condition are shown. Subjective day and night are depicted by the gray and black bars, respectively. (B) Circadian period lengths for +, *chico*^*1*^ / + (BC5), *chico*^*1*^ / + (F1), and *chico*^*1*^ flies. Circadian period lengths were calculated using locomotor activity data for 10 days by FT. (C) Amplitudes of circadian rhythm for +, *chico*^*1*^ / + (BC5), *chico*^*1*^ / + (F1), and *chico*^*1*^. Amplitudes were calculated using locomotor activity data for 10 days by FT. *** p < 0.001; *t*-test.

### Insulin signaling in neurons, especially in the clock neurons, regulates circadian rhythm

To determine whether the neuronal insulin signaling regulates circadian rhythm, we inhibited the insulin signaling in specific regions using Gal4 / upstream activation sequence (UAS) system (Brand and Perrimon, 1993). First, we used pan-neuronal Gal4 driver, *elav*-Gal4, and UAS-*chico* RNAi to obtain the flies with pan-neuronal knockdown of *chico*. The results showed that the circadian period length was significantly shortened in flies with pan-neuronal knockdown of *chico* (Fig. 2A, Sup. Fig. 1A). Also, we conducted the expression of a dominant-negative form of *InR* (*InR* DN) and knockdown of *InR* or *dp110* (*Phosphatidylinositol 3-kinase 92E*), one of the downstream of insulin signaling, to validate the effect of insulin signaling inhibition on circadian rhythm. These manipulations with *elav*-Gal4 caused shorter circadian period length as well as pan-neuronal knockdown of *chico* (Fig. 2A, Sup. Fig. 1B-D). In addition, pan-neuronal knockdown of *InR* caused decreased amplitude of circadian rhythm (Fig. 2B). To confirm the efficiency of RNAi, we quantified the mRNA level of each gene and *4E-BP*, the expression of which is negatively regulated by insulin signaling (Jünger et al., 2003; Puig et al., 2003). Although *4E-BP* mRNA level was significantly increased only by the pan-neuronal knockdown of *chico*, the mRNA level of *chico* or *dp110* was significantly declined in flies with pan-neuronal knockdown of *chico* or *dp110*, respectively (Sup. Fig. 2A, C). Although we couldn’t detect any change of *InR* mRNA level in flies with pan-neuronal knockdown of *InR*, there is a significant increase of *4E-BP* mRNA level in these flies (Sup. Fig. 2B, D), suggesting the inhibition of insulin signaling. Taken together, insulin signaling in the brain could regulate circadian rhythm.

**Figure 2.**
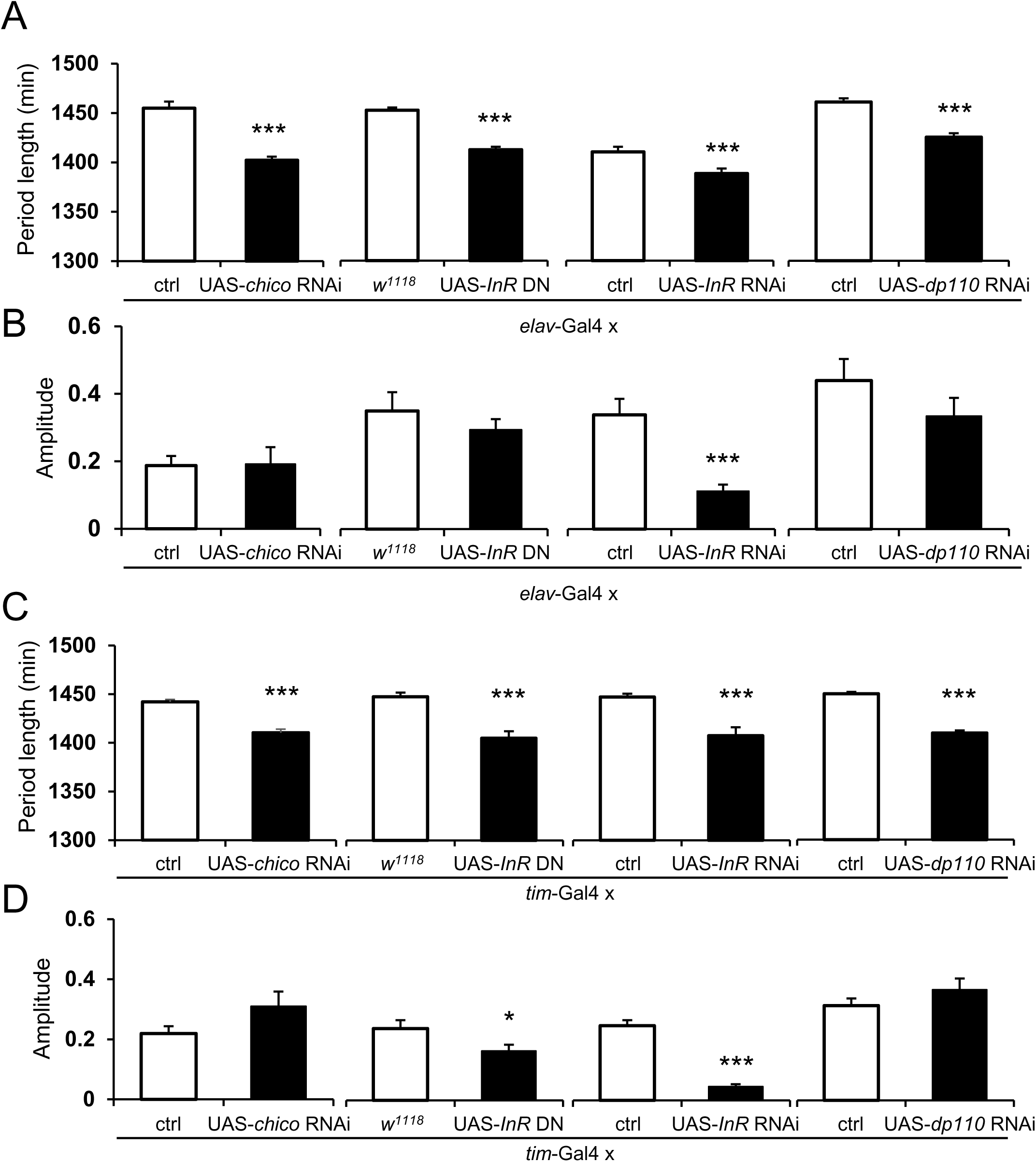
Inhibition of insulin signaling in clock neurons resulted in shorter circadian period length. (A, B) Circadian period lengths (A) and amplitudes of circadian rhythm (B), which were calculated using locomotor activity data for 10 days by FT, for each genotype denoted at bottom of the panels. *w*^*1118*^, the genetical background of UAS-*InR* DN, was used as a control for UAS-*InR* DN. *y*^*1*^ *v*^*1*^; *P{CaryP}attP2* was used as ctrl for UAS-*chico* RNAi. *y*^*1*^ *v*^*1*^; *P{CaryP}attP40* was used as ctrl for UAS-*InR* RNAi and UAS-*dp110* RNAi. Since each experiment was conducted independently, the control group for UAS-*InR* RNAi and UAS-*dp110* RNAi showed different circadian period lengths and amplitudes, although they are the same genotype. The numbers of flies of each bar were 31, 31, 29, 32, 16, 16, 16, and 11 from the left. (C, D) Circadian period lengths (C) and amplitudes of circadian rhythm (D), which were calculated using locomotor activity data for 10 days by FT, for each genotype denoted at bottom of the panels. The numbers of flies of each bar were 32, 27, 32, 32, 32, 32, 32, and 32 from the left. * p < 0.05, *** p < 0.001; *t*-test.

Since inhibition of neuronal insulin signaling affected behavioral rhythm, we focused on the clock neurons, which express clock genes and are essential for the generation of circadian rhythms. (Dubowy and Sehgal, 2017). To manipulate the insulin signaling in the clock neurons, we employed *tim*-Gal4 which expresses GAL4 under the promoter of *timeless*, one of the clock genes in *Drosophila* (Myers et al., 1995; Sehgal et al., 1995). Inhibition of insulin signaling with *tim*-Gal4 shortened circadian period length (Fig. 2C, Sup. Fig. 3A-D). Moreover, the expression of *InR* DN or knockdown of *InR* declined the amplitudes of the circadian rhythm (Fig. 2D). These results indicated that insulin signaling in clock neurons regulates circadian rhythm.

**Figure 3.**
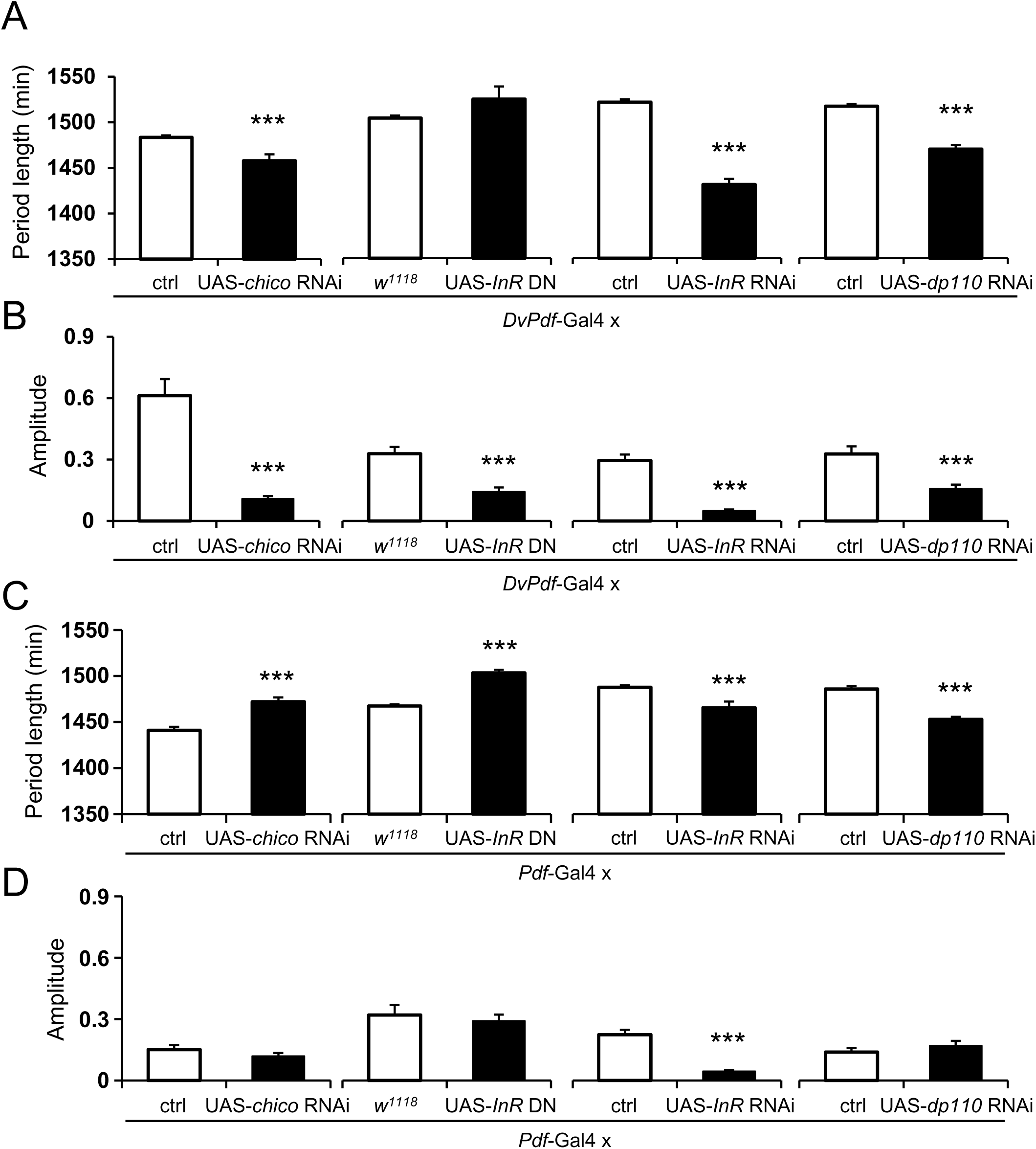
Insulin signaling in the lateral neurons regulates circadian rhythm. (A, B) Circadian period lengths (A) and amplitudes of circadian rhythm (B), which were calculated using locomotor activity data for 10 days by FT, for each genotype denoted at bottom of the panels. *w*^*1118*^, the genetical background of UAS-*InR* DN, was used as a control for UAS-*InR* DN. *y*^*1*^ *v*^*1*^; *P{CaryP}attP2* was used as ctrl for UAS-*chico* RNAi. *y*^*1*^ *v*^*1*^; *P{CaryP}attP40* was used as ctrl for UAS-*InR* RNAi and UAS-*dp110* RNAi. Since each experiment was conducted independently, the control group for UAS-*InR* RNAi and UAS-*dp110* RNAi showed different circadian period lengths and amplitudes, although they are the same genotype. The numbers of flies of each bar were 32, 31, 31, 30, 32, 32, 32, and 32 from the left. (C, D) Circadian period lengths (C) and amplitudes of circadian rhythm (D), which were calculated using locomotor activity data for 10 days by FT, for each genotype denoted at bottom of the panels. The numbers of flies of each bar were 32, 32, 32, 31, 32, 32, 29, and 28 from the left. Error bars indicate SEM. *** p < 0.001; *t*-test.

Among the clock neurons, ventral and dorsal lateral neurons (LNvs and LNds) are recently demonstrated to be the two key components of the clock neuron network (Delventhal et al., 2019; Schlichting et al., 2019). These studies showed that the lack of a clock gene both in LNvs and LNds caused complete loss of behavioral rhythmicity and the molecular clock either in LNvs or in LNds is sufficient to generate the behavioral circadian rhythm. On the other hand, LNvs themselves are known to be necessary to synchronize the clock neuron network via the release of a principal circadian messenger, pigment dispersing factor (PDF) (Renn et al., 1999). To validate whether insulin signaling both in LNvs and LNds or only in LNvs relates to circadian rhythm regulation, inhibition of insulin signaling was conducted by *DvPdf*-Gal4 and *Pdf*-Gal4. As a result, knockdown of *chico, InR* or *dp110* by *DvPdf*-Gal4 shortened circadian period length (Fig. 3A, Sup. Fig. 4A-D). Although expression of *InR* DN with *DvPdf*-Gal4 did not significantly change period length, it caused arrhythmic-like behavior after four to five days of DD conditions (Sup. Fig. 4B). Furthermore, all these manipulations with *DvPdf*-Gal4 decreased the amplitudes of the circadian rhythm (Fig. 3B). In addition, the knockdown of *InR* or *dp110* with *Pdf*-Gal4 shortened circadian period length (Fig. 3C, Sup. Fig. 5A-D). The *InR* knockdown by *Pdf*-Gal4 weakened the circadian rhythm amplitude (Fig. 3D). On the other hand, both the knockdown of *chico* and the expression of *InR* DN with *Pdf*-Gal4 caused a longer circadian period length (Fig. 3C). These results indicated that insulin signaling in LNvs contribute to maintain the proper behavioral rhythm.

**Figure 4.**
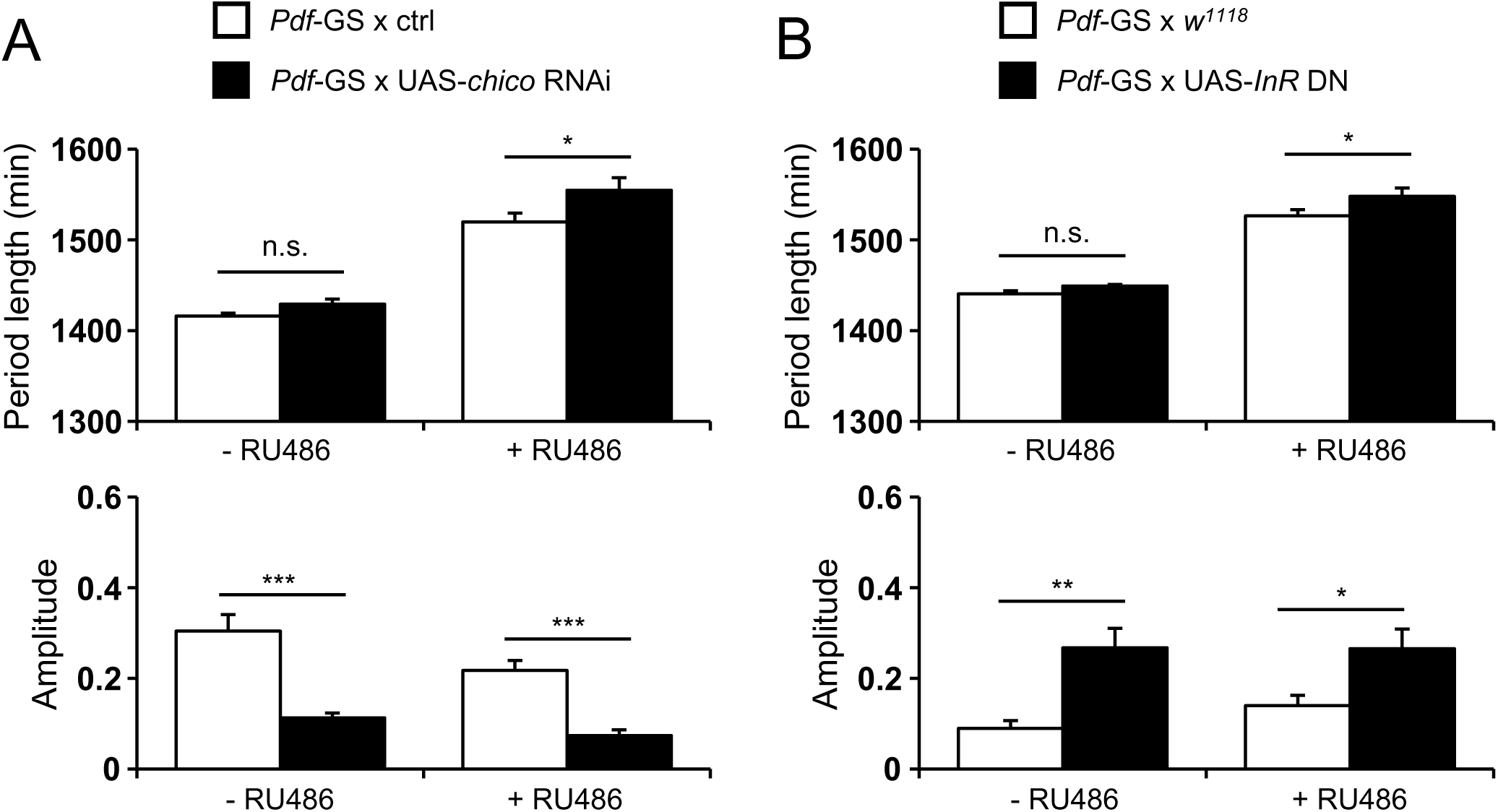
Insulin signaling in LNvs regulates circadian rhythm in a development-independent manner. (A) Circadian period lengths and amplitudes of circadian rhythm, which were calculated using locomotor activity data for 10 days by FT, for control (*Pdf*-GS x ctrl, white bar) and flies with *chico* knockdown by *Pdf*-GS (*Pdf*-GS x UAS-*chico* RNAi, black bar) without RU486 (- RU486) or with RU486 (+ RU486). *y*^*1*^ *v*^*1*^; *P{CaryP}attP2* was used as ctrl for UAS-*chico* RNAi. The numbers of flies of each bar were 24, 28, 18, and 20 from the left. Error bars indica te SEM. (B) Circadian period lengths and amplitudes of circadian rhythm, which were calculated using locomotor activity data for 10 days by FT, for control (*Pdf*-GS x *w*^*1118*^, white bar) and flies with *InR* DN expression by *Pdf*-GS (*Pdf*-GS x UAS-*InR* DN, black bar) without RU486 (- RU486) or with RU486 (+ RU486). The numbers of flies of each bar were 26, 25, 27, and 26 from the left. Error bars indicate SEM. *w*^*1118*^, the genetical background of UAS-*InR* DN, was used as a control for UAS-*InR* DN. n.s. p ≥ 0.05, * p < 0.05, ** p < 0.01, *** p < 0.001; Tukey Kramer method.

**Figure 5.**
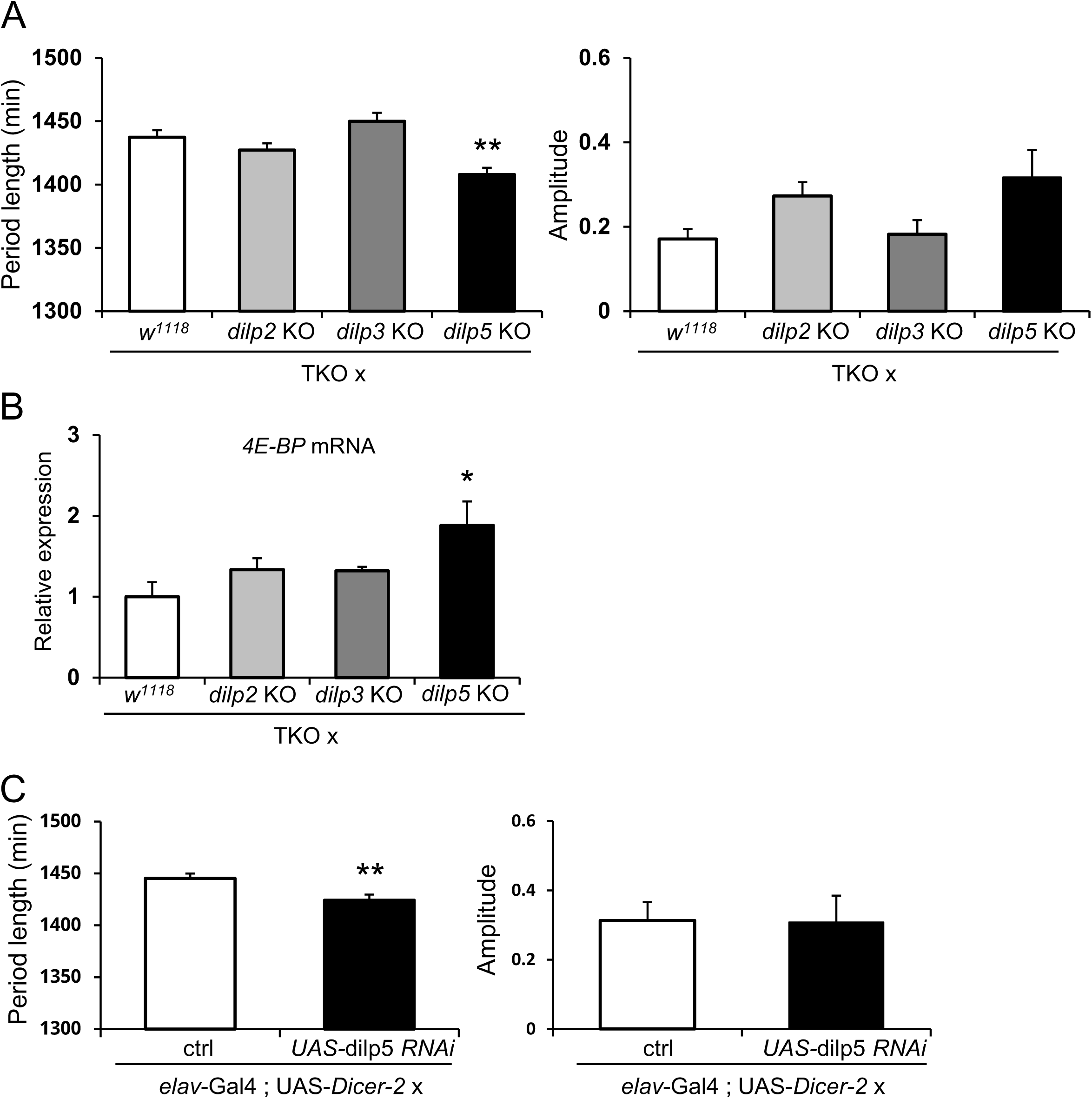
*dilp5* relates to the regulation of circadian rhythm by insulin signaling. (A) Circadian period lengths and amplitudes of circadian rhythm, which were calculated using locomotor activity data for 10 days by FT, for each genotype denoted at bottom of the panels: TKO x *w*^*1118*^ (n = 15), TKO x *dilp2* KO (n = 16), TKO x *dilp3* KO (n = 16), TKO x *dilp5* KO (n = 13). (B) Relative expression of *4E-BP* for each genotype denoted at bottom of each panel. (C) Circadian period lengths and amplitudes of circadian rhythm, which were calculated using locomotor activity data for 10 days by FT, for *elav*-Gal4; UAS-*Dicer-2* x ctrl (n = 14) and *elav*-Gal4; UAS-*Dicer-2* x UAS-*dilp5* RNAi (n = 11). *w*^*1118*^(VDRC stock number: 60000) was used as ctrl for UAS-*dilp5* RNAi. Error bars indicate SEM. * p < 0.05, ** p < 0.01; vs. TKO x *w*^*1118*^; Tukey Kramer method (A, B). ** p < 0.01; *t*-test (C)

Since insulin signaling relates to development and growth (Naganos et al., 2012; Song et al., 2003), constitutive inhibition of insulin signaling could cause behavioral changes via some developmental defects. To avoid inhibition of insulin signaling during development and growth stages, we used a drug-inducible gene expression system, Gene-Switch (GS) system (McGuire et al., 2004). Knockdown of *chico* and expression of *InR* DN specifically at the adult stage were achieved by using *Pdf*-GS. Without RU486, flies with UAS-*chico* RNAi or UAS-*InR* DN did not show significant changes in circadian period length (Fig. 4A, B, Sup. Fig. 6A, B). With RU486, flies with *chico* knockdown or expression of *InR* DN by *Pdf*-GS showed significantly longer circadian period length (Fig. 4A, B). Since the amplitudes from flies with UAS-*chico* RNAi or UAS-*InR* DN are different from each control without RU486, there might be some genetical background effects. These results indicated that insulin signaling in LNvs relates to circadian rhythm regulation, at least in part, independently of its developmental function.

### *dilp5* contributes to the regulation of circadian rhythm via insulin signaling

Next, we focused on *dilp2, dilp3*, and *dilp5*, which are expressed in the brain, among the eight *dilps* (Nässel et al., 2015; Nässel and Broeck, 2016). To inhibit the compensational expression of other *dilps* in a knockout mutant of one *dilp* (Grönke et al., 2010), we crossed every single knockout mutant of *dilp2* (*dilp2* KO), *dilp3* (*dilp3* KO), or *dilp5* (dilp5 KO) with a triple knockout mutant of *dilp2, dilp3*, and *dilp5* (TKO) and analyzed the flies of one *dilp* homozygous KO with two heterozygous KO. Among these mutants, only *dilp5* KO (TKO x *dilp5* KO) showed a significantly shorter circadian period length than control (TKO crossed with *w*^*1118*^, TKO x *w*^*1118*^) (Fig. 5A, Sup. Fig. 7). As we previously reported, each heterozygous mutant expressed an excess amount of *dilp* other than systematically knockout one (Yamaguchi et al., 2021). To elucidate the effect of systematically *dilps* knockout on insulin signaling, the mRNA level of *4E-BP* was quantified. *4E-BP* expression was significantly increased only in TKO x *dilp5* KO, suggesting that insulin signaling is inhibited in TKO x *dilp5* KO (Fig. 4B). On the other hand, *4E-BP* mRNA level did not change in the heterozygous mutant of TKO and *dilp2* KO (TKO x *dilp2* KO) or *dilp3* KO (TKO x *dilp3* KO), despite their mutation in *dilps*. To confirm the function of *dilp5* in the regulation of circadian rhythm, we knocked down *dilp5* with *elav*-Gal4. Accordingly, flies with pan-neuronal knockdown of *dilp5* showed a significant reduction of circadian period length without any changes of mRNA level of *dilp2* or *dilp3* (Fig. 5C, Sup. Fig. 8). These results indicated the regulation of circadian rhythm by *dilp5*.

## Discussion

In this study, we investigated the relationship between circadian rhythm and insulin signaling. To clarify the relationship, we conducted inhibition of insulin signaling in the targeted region. We also analyzed the activity of the *dilps* mutants to specify the ligand involved in the regulation of circadian rhythm. Through these experiments, we showed that *InR* and its downstream molecules in the clock neurons regulate circadian rhythm. In addition, we discovered that *dilp5* contributes to the regulation of circadian rhythm. These findings support the relationship between circadian rhythm and metabolism in *Drosophila melanogaster*.

Specifically, we examined the effect of inhibition of insulin signaling on circadian rhythm by the expression of a dominant-negative form of *InR, InR* DN, and by the knockdown of *chico, InR*, and *dp110*. All of these manipulations with *elav-*Gal4 and *tim*-Gal4 drivers shortened circadian period length (Fig. 2A, Fig. 2C), although we couldn’t detect a significant change of *InR* expression in flies with pan-neuronal knockdown of *InR*. It may be because *InR* is expressed in various tissues, including the neurons, glia, and fat body. Knockdown of the three genes with *DvPdf*-Gal4 also caused the same phenotype in period length (Fig. 3A). Also, knockdown of *InR* or *dp110* by *Pdf*-Gal4 shortened circadian period length. On the other hand, knockdown of *chico* or expression of *InR* DN by *Pdf*-Gal4 lengthened period length (Fig. 3C). In previous research, flies with silencing of *TOR* in the cells expressing *period*, one of the core clock genes, showed the altered circadian period length (Kijak and Pyza, 2017). Other paper reported that activation of *Akt* and *TOR* signaling in LNvs lengthened circadian period length(Zheng and Sehgal, 2010). Although the detailed distribution of *InR* is poorly understood to date, cell-specific RNA sequencing of each clock cell detected mRNA expression of the insulin signaling molecules in the clock neurons (Abruzzi et al., 2017; Ma et al., 2021). These results support the idea that neuronal insulin signaling, especially in the clock neurons, regulates circadian rhythm.

In our experiment, most of the neuronal insulin signaling inhibition altered circadian period length. Although only the expression of *InR* DN with *DvPdf*-Gal4 did not cause significant changes in the circadian period length (Fig. 3A), these flies seem to have a shorter circadian period length than control according to the activity in the first 4-days of DD condition (Sup. Fig. 4B). Also, these flies showed arrhythmic-like behavior after the 4-5 days of DD condition. Such phenotype might cause a loss of accuracy for quantification of circadian period length. Contrary to the shortened circadian period length in the knockdown of *InR* or *dp110* by *Pdf*-Gal4, the knockdown of *chico* and the expression of *InR* DN with *Pdf*-Gal4 lengthened circadian period length (Fig. 3C). There are some possible causes of these contradictions, such as the genetic background, the strength of insulin signaling inhibition, or the developmental defects. Although there were some contradictions, we speculated that the insulin signaling in the clock neurons contribute to maintain the proper behavioral rhythm.

Recently, Schlichting et al. reported that knockout of *period* in all or subset of lateral neurons differently affected circadian period length. In addition, Schlichting et al. and Delventhal et al. demonstrated that the clock neurons act not in a hierarchy but as a distributed network to regulate circadian activity (Delventhal et al., 2019; Schlichting et al., 2019). In these papers, cell-specific knockout of clock genes revealed that the clock in each clock neuron, especially in LNvs and LNds, collaborate to achieve the close to 24-hr period length. Inhibition of insulin signaling in the lateral neurons might diminish such communication between the lateral neurons. Although inhibition of insulin signaling by *elav*-Gal4 and *tim*-Gal4 did not cause severe defects in circadian behavior as much as *DvPdf*-Gal4 did, it might be due to the difference of Gal4 expression levels in LNvs and LNds. Furthermore, *shaggy* (*sgg*, homolog of *GSK3*), which is negatively regulated by insulin signaling (Papadopoulou et al., 2004), is known to phosphorylate clock protein and manipulate the intrinsic speed of the molecular clock (Martinek et al., 2001). Additionally, Zhang and Sehgal reported that *Akt* and *TOR* modulate the timing of nuclear expression of TIM in the clock neurons (Zheng and Sehgal, 2010). Although we did not assess whether our manipulation of insulin signaling changes the core clock gene behavior or neural communication, it is possible that insulin signaling modulated the molecular clock via the regulation of *sgg* activity as previous research reported. Further experiments are needed to clarify the effect of insulin signaling on the clock gene expression rhythm or the clock neuron activity.

Since insulin signaling is important for development, the inhibition of insulin signaling with constitutive Gal4 might induce some developmental defects (Böhni et al., 1999; Naganos et al., 2012; Song et al., 2003). However, we demonstrated that the adult stage-specific insulin signaling inhibition in LNvs by *Pdf*-GS altered behavioral rhythm (Fig. 4A, B). These results suggested that insulin signaling, at least in LNvs, regulates circadian rhythm in a development-independent manner.

We demonstrated that *dilp5* alone among three insulin ligands tested showed a significant effect on the circadian period length when only one of the *dilps* was systemically knocked out or knocked down in neurons. An earlier study reported that *dilp* single mutation caused compensational expression of other *dilps* and little change of *4E-BP* expression (Grönke et al., 2010). We previously reported the similar compensational effects in the progenies from crosses between TKO and single KO mutants (Yamaguchi et al., 2021). Especially, *dilp2* mRNA level in the mutants of *dilp5* was significantly increased. *dilp2* mRNA increase was also observed in the mutants of *dilp3* which did not show any changes in circadian period length. In addition, pan-neuronal knockdown of *dilp5* shortened circadian period length without an increase of other *dilps* mRNA (Fig. 5F, Sup. Fig. 8B). According to these results, we conjectured that increase of *dilp2* mRNA did not cause the changes of circadian period length. However, it is still possible that *dilp2* or *dilp3* plays a role in the regulation of circadian rhythm. It should also be noted that *dilps*, especially *dilp2, dilp3*, and *dilp5*, are thought to have a functional redundancy (Nässel et al., 2015). Nevertheless, we demonstrated that *dilp5* has a significant effect on circadian rhythm.

In conclusion, our results suggest that neuronal insulin signaling contributes to the regulation of robust circadian rhythm in flies. In mammals, previous studies showed the relationship between insulin signaling and circadian rhythm (Boden et al., 1996; Marcheva et al., 2010; Sato et al., 2014). Recently, Crosby et al. demonstrated that insulin and insulin-like growth factor 1 are important for the entrainment of behavioral rhythms to feeding time and have some activity in the suprachiasmatic nucleus, the master clock of mammals (Crosby et al., 2019). Taken together, our results shed light on the conserved role of insulin and insulin signaling in the regulation of circadian rhythm from flies to mammals.

## Acknowledgments

We thank Dr. Taishi Yoshii, BDSC, and VDRC for fly stocks, and the members of Kume lab for discussions.

