## Supplemental Figures for "The regulation of circadian rhythm by insulin signaling in *Drosophila*"

**Supplemental Figure Legends**

**Supplemental Figure 1. Average activity of flies with pan-neuronal insulin signaling inhibition**

(A-D) Average activity from flies with pan-neuronal expression of *chico* RNAi (A), *InR* DN (B), *InR* RNAi (C), or *dp110* RNAi(D), and each left panel shows the average activity from control. Each fly activity was recorded for 10 days in constant darkness (DD) conditions. Activity counts over 7 days in DD condition is shown. Subjective day and night are depicted by the gray and black bars, respectively.

**Supplemental Figure 2. Knockdown efficacy of *chico*, *InR*, or *dp110*.**

(A-D) relative expression of *chico* (A), *InR* (B), *dp110* (C), and *4E-BP* (D) for each genotype denoted at bottom of panels. Error bars indicate SEM. *p < 0.05 vs. *elav*-Gal4 x ctrl; Tukey-Kramer method (D, for comparison between *elav*-Gal4 x ctrl, *elav*-Gal4 x UAS-*InR* RNAi, and *elav*-Gal4 x UAS-*dp110* RNAi). * p < 0.05, ** p < 0.01; *t*-test (A-D).

**Supplemental Figure 3. Average activity of flies with insulin signaling inhibition in cells expressing the clock gene.**

(A-D) Average activity from flies expressing *chico* RNAi (A), *InR* DN (B), *InR* RNAi (C), or *dp110* RNAi(D) with *tim*-Gal4, and each left panel shows the average activity from control. Each fly activity was recorded for 10 days in DD conditions. Activity counts over 7 days in DD condition is shown. Subjective day and night are depicted by the gray and black bars, respectively.

**Supplemental Figure 4. Average activity of flies with inhibition of insulin signaling in LNvs and LNds.**

(A-D) Average activity from flies expressing *chico* RNAi (A), *InR* DN (B), *InR* RNAi (C), or *dp110* RNAi(D) with *DvPDF*-Gal4, and each left panel shows the average activity from control. Each fly activity was recorded for 10 days in DD conditions. Activity counts over 7 days in DD condition is shown. Subjective day and night are depicted by the gray and black bars, respectively.

**Supplemental Figure 5. Average activity of flies with inhibition of insulin signaling in LNvs.**

(A-D) Average activity from flies expressing *chico* RNAi (A), *InR* DN (B), *InR* RNAi (C), or *dp110* RNAi(D) with *PDF*-Gal4, and each left panel shows the average activity from control. Each fly activity was recorded for 10 days in DD conditions. Activity counts over 7 days in DD condition is shown. Subjective day and night are depicted by the gray and black bars, respectively.

**Supplemental Figure 6. Average activity of flies with inhibition of insulin signaling in LNvs specifically at adult stage.**

(A, B) Average activity from flies of each genotype (control for *chico* RNAi: *Pdf*-GS x *ctrl*, flies with chico RNAi: *Pdf*-GS x UAS-*chico* RNAi, control for *InR* DN: *Pdf*-GS x $w^{1118}$, flies with *InR* DN: *Pdf*-GS x UAS-*InR* DN) without RU486 (- RU486) or with RU486 (+ RU486) denoted at each top of panel. Each fly activity was recorded for 10 days in DD conditions. Activity counts over 7 days in DD condition is shown. Subjective day and night are depicted by the gray and black bars, respectively.

**Supplemental Figure 7. Average activity of flies with mutation in *dilps*.**

Average activity from flies of each genotype denoted at top of each panel. Each fly activity was recorded for 10 days in DD conditions. Activity counts over 7 days in DD condition is shown. Subjective day and night are depicted by the gray and black bars, respectively.

**Supplemental Figure 8. Average activity of flies with pan-neuronal knockdown of *dilp5*.**

(A) Average activity from flies expressing *dilp5* RNAi (A) with *elav*-Gal4, and left panel shows the average activity from control. Each fly activity was recorded for 10 days in DD conditions. Activity counts over 7 days in DD condition is shown. Subjective day and night are depicted by the gray and black bars, respectively. (B) Relative expression of *dilp2*, *dilp3*, or *dilp5* for control and flies with pan-neuronal knockdown of *dilp5*. Error bars indicate SEM. *** $p < 0.001$; *t*-test.

### Supplemental Figure 1

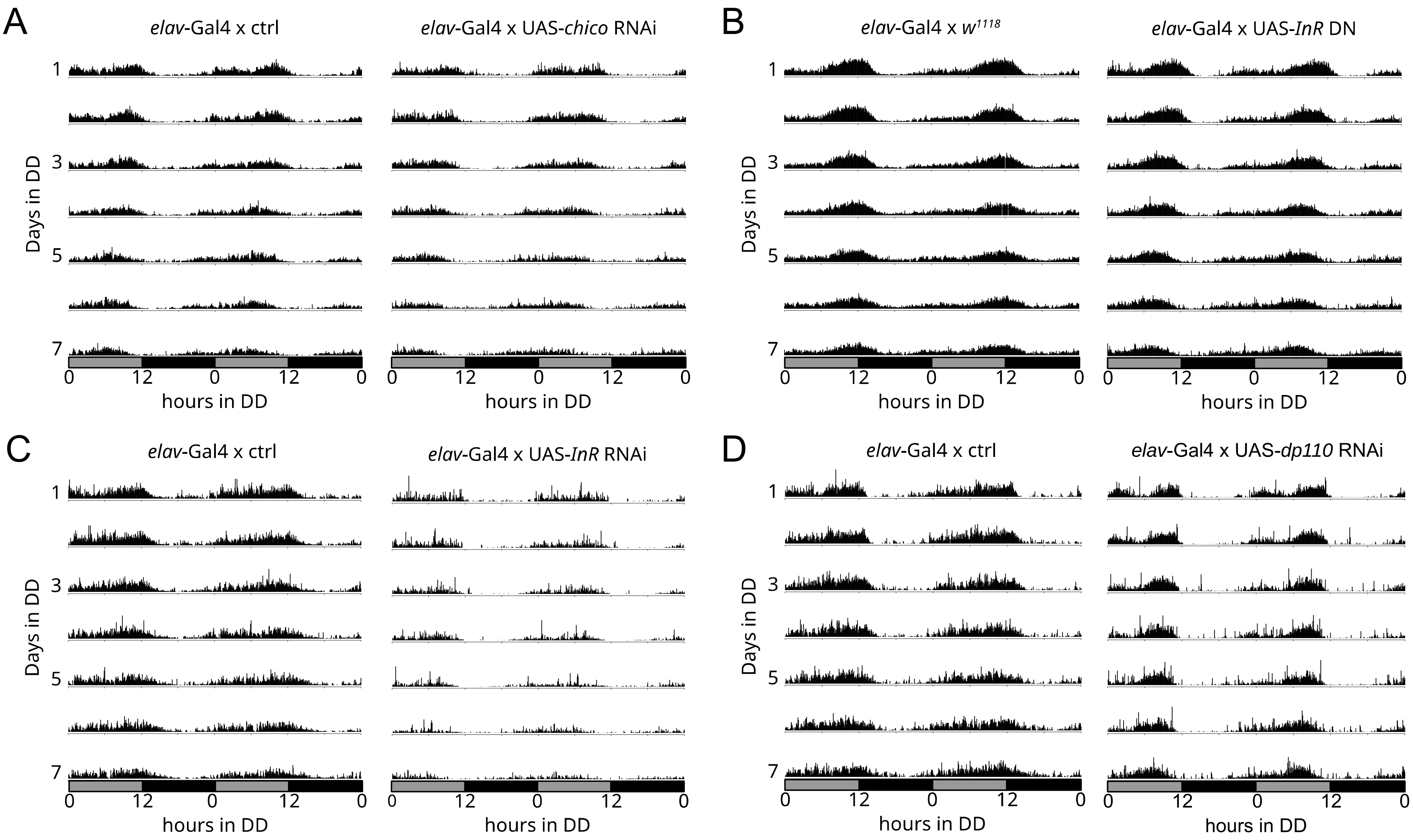

### Supplemental Figure 2

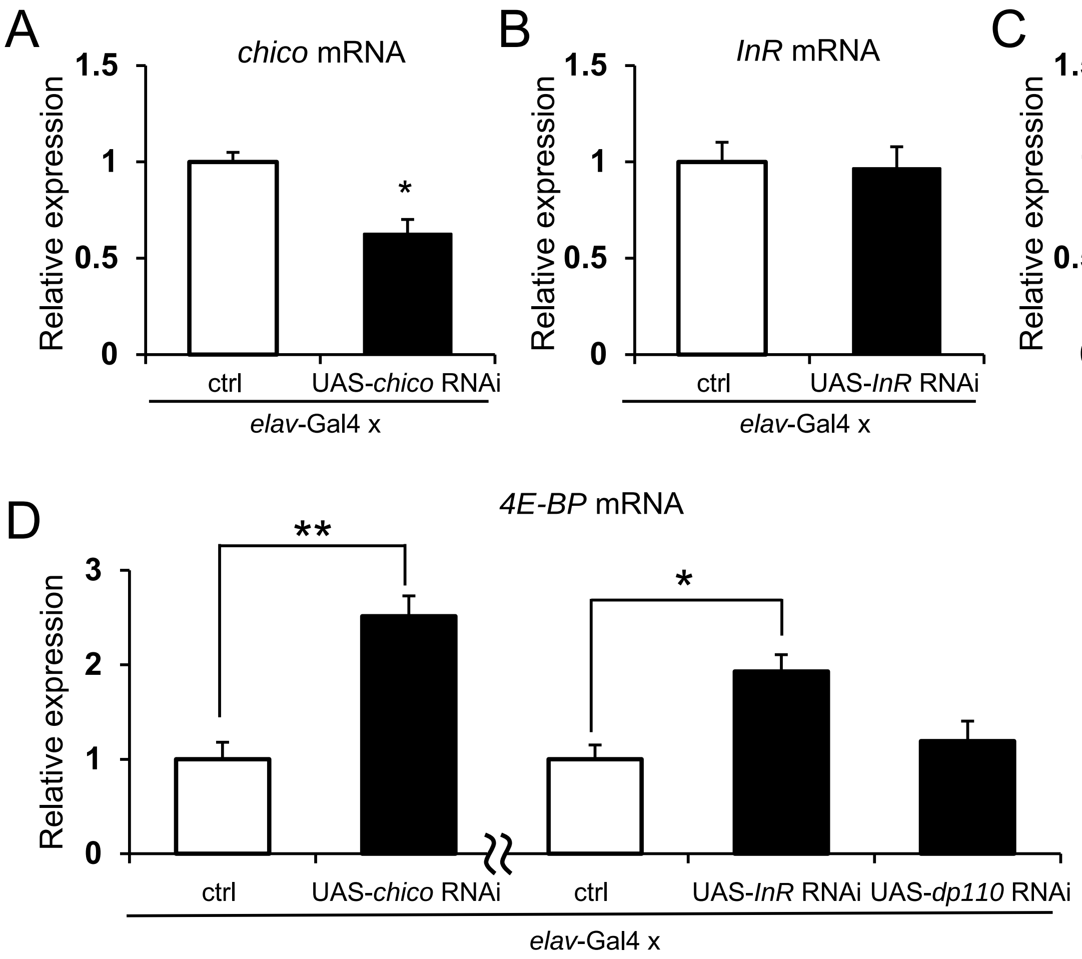

### Supplemental Figure 3

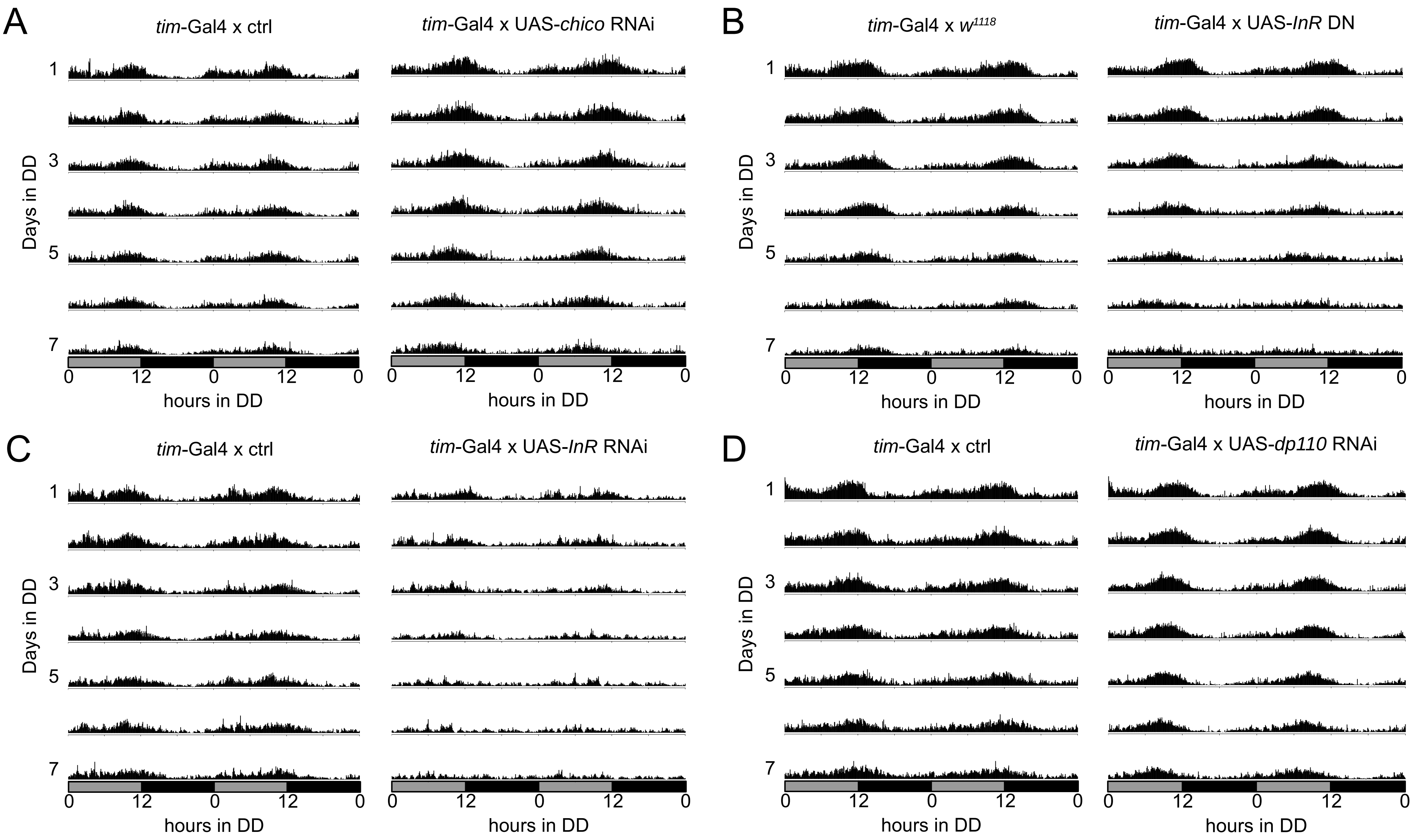

### Supplemental Figure 4

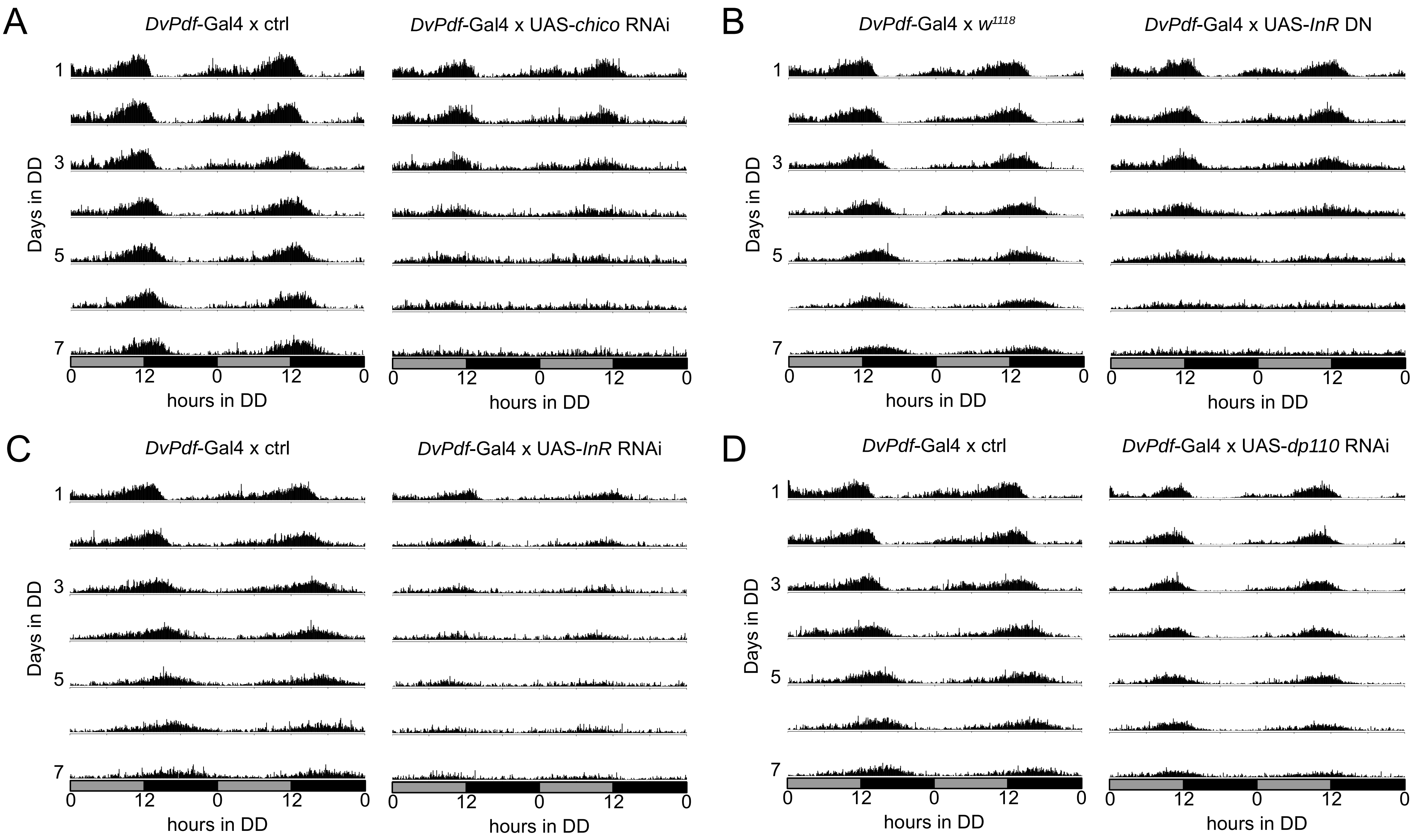

### Supplemental Figure 5

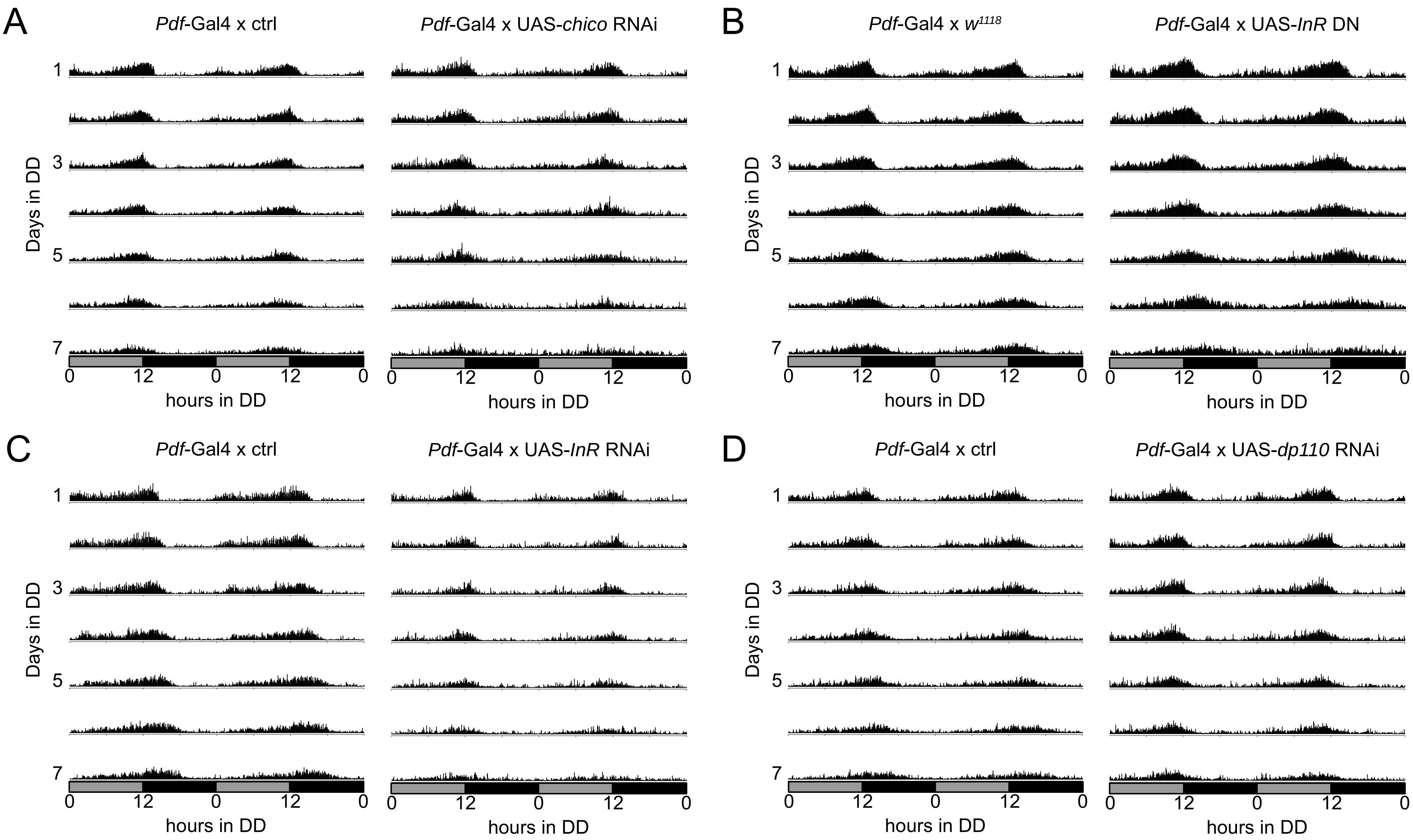

### Supplemental Figure 6

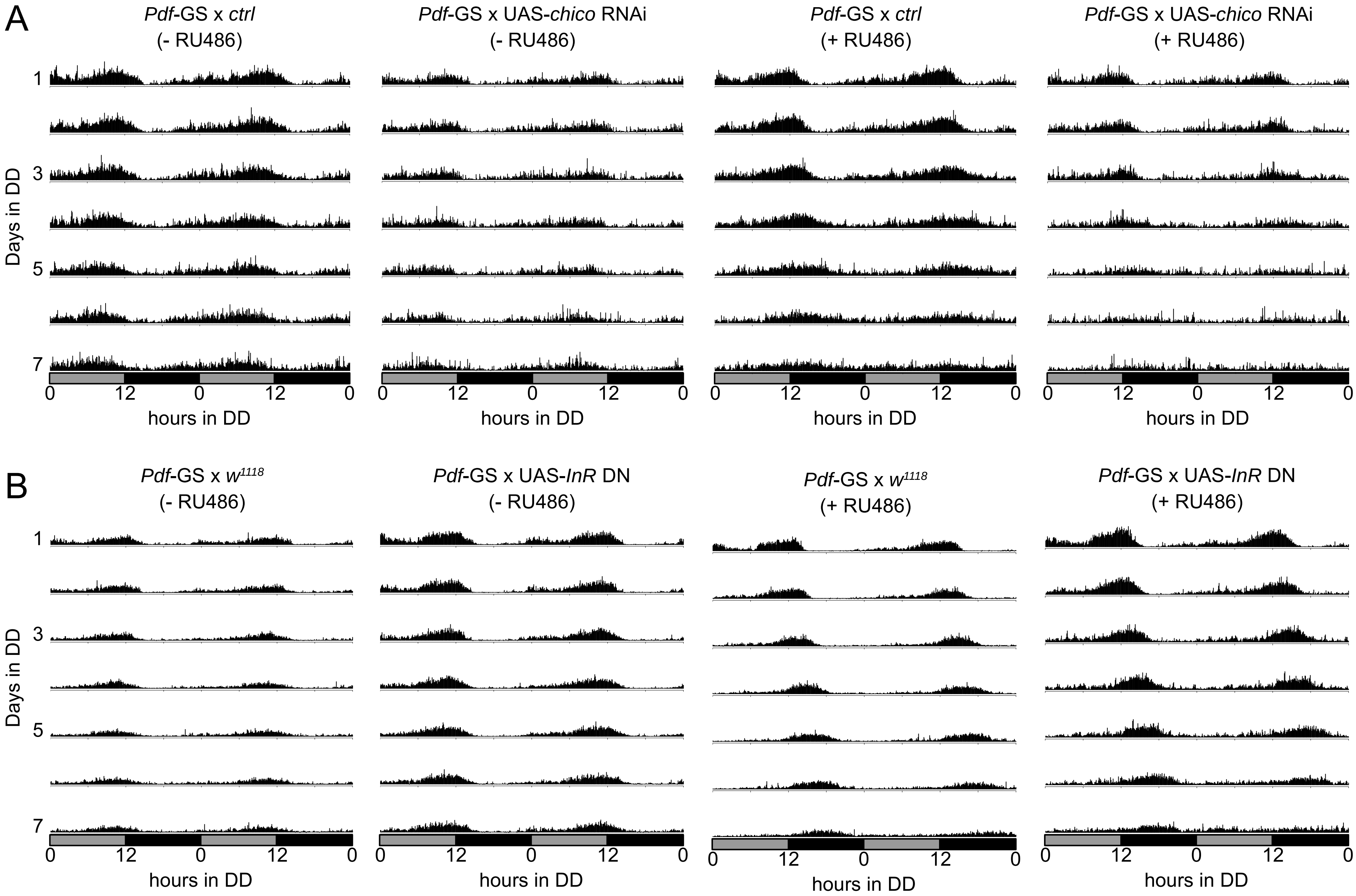

### Supplemental Figure 7

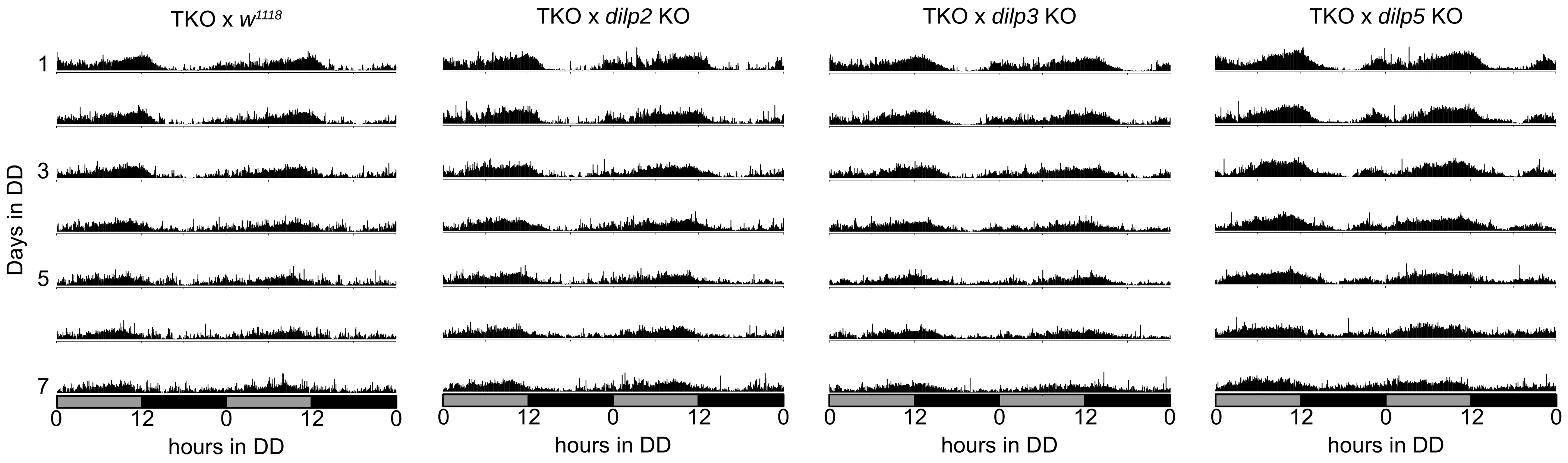

### Supplemental Figure 8

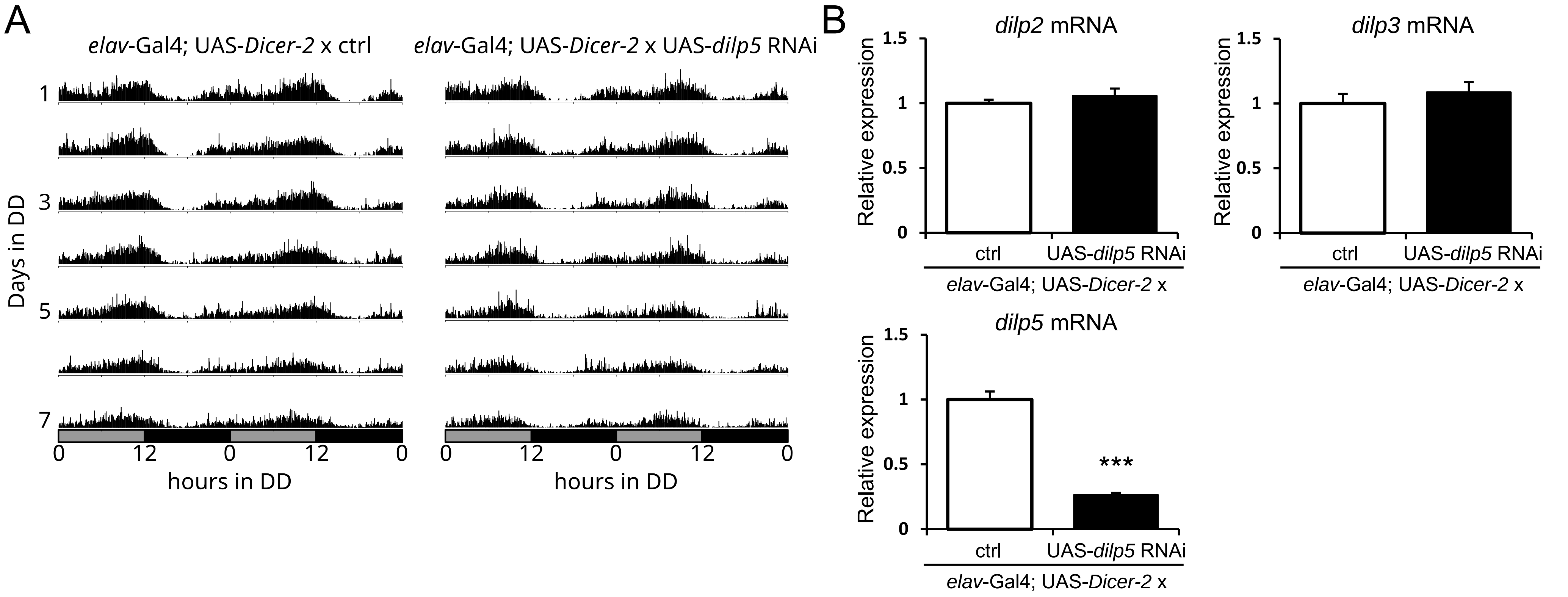
